# Circulating CD4 regulatory T cells are broadly preserved in Parkinson’s disease despite altered HELIOS expression

**DOI:** 10.64898/2026.11.06.764595

**Authors:** Sara C. Crooks, Aaditya Prabhu, Lorna B. Jarvis, Zoya G. Georgieva, Daniel B. Rainbow, Caroline H. Williams-Gray, Joanne L. Jones

## Abstract

**Background:** Regulatory T cells (Tregs) are an important component of adaptive immune regulation and have been implicated in Parkinson’s disease (PD), with previous studies suggesting altered Treg proportions, phenotype and, in some cases, function. However, existing studies have been limited by variable phenotyping approaches and a lack of detailed functional assessment. Given the growing interest in Tregs as potential therapies in PD, better understanding of their phenotype and function in the disease is important.

**Methods:** Circulating FOXP3+ Tregs were characterised in individuals with PD and age-matched healthy controls (HC) using a 24-colour spectral flow cytometry panel. Treg suppressive function was assessed by their ability to inhibit proliferation and cytokine production by polyclonally stimulated autologous T effector cells *in vitro*. Treg lineage stability was assessed by analysis of *FOXP3* TSDR demethylation.

**Results:** The proportions and phenotypic characteristics of circulating CD4 Tregs were broadly similar between individuals with PD and HCs, as were Treg suppressive function and *FOXP3* TSDR demethylation. However, naive CD4 Tregs from individuals with PD showed reduced HELIOS expression, with a similar but non-significant trend in FOXP3 expression. Reduced HELIOS expression was also observed in the CD8 naive Treg compartment.

**Conclusions:** Major quantitative or functional abnormalities in circulating Tregs do not appear to be a dominant feature PD. However, reduced HELIOS expression in naive Tregs suggests more subtle phenotypic alterations that warrant further investigation. The broadly preserved suppressive function is reassuring for therapeutic approaches aimed at increasing Treg numbers, although it will be important to establish that expanded Tregs retain a stable phenotype and suppressive function in inflammatory and tissue environments.

## 1. Introduction

Inflammation is increasingly recognised as a pivotal feature of Parkinson’s disease (PD) pathology. Beyond the central nervous system, there is increasing interest in the role of the peripheral innate and adaptive immune responses in PD, including monocytes, B cells and NK cells as well as T cells(Bhatia et al., 2021; Holbrook et al., 2023; Li et al., 2021; Sun et al., 2019; Wijeyekoon et al., 2020). Genetic studies have implicated the HLA Class II region in PD susceptibility. In particular, the intronic variant (rs3129882) in *HLA-DRA* is associated with late-onset and sporadic PD and has been shown to alter the expression of HLA Class II genes (Hamza et al., 2010). A more recent study found four HLA types associated with PD and that certain *HLA-DRB* subtypes are protective and associated with lower risk of PD, which was supported in a mouse model(Carver et al., 2026; Yu et al., 2021). Epidemiological studies have also associated regular use of non-steroidal anti-inflammatories (NSAIDs) (not including aspirin) with a lower risk of PD with stronger associations observed with longer durations of use(H. Chen et al., 2003). NSAID use has also been associated with lower risk of PD in people with *LRRK2* mutations(Luciano et al., 2020), suggesting that reducing systemic inflammation might be beneficial in reducing the risk of developing PD.

Multiple studies have suggested alterations in regulatory T cells (Tregs), a small subset (5-10%) of CD4 T cells that play an important role in maintaining immune homeostasis and preventing excessive immune activation(Fehérvari & Sakaguchi, 2004; Halim et al., 2017). However, evidence for quantitative and functional alterations in Tregs in PD remains inconsistent. Several studies have reported lower proportions of Tregs in PD(Y. Chen et al., 2015; Thome et al., 2021; Yang et al., 2016.), including reductions in what they define as active (CD45RO+) Tregs(Álvarez-Luquín et al., 2019). In contrast, others have found no differences in Treg frequency between individuals with PD and controls(Cen et al., 2017; Karaaslan et al., 2021; Saunders et al., 2012). Importantly, Treg definitions have varied between studies, with some including FOXP3 expression, the lineage-defining transcription factor, and others relying purely on expression of surface markers such as CD4, CD25 and CD127.

Functional studies have similarly provided mixed evidence, with some reporting reduced suppressive capacity, although not consistently across different Teffector:Treg ratios(Saunders et al., 2012; Thome et al., 2021). More recently, reduced FOXP3 mean fluorescence intensity (MFI; indicating density of expression) of CD4 Tregs (CD127lowCD25highFOXP3+) and reduced HELIOS MFI of HELIOS+ CD4 Tregs (CD127lowCD25highFOXP3+HELIOS+) was reported in a cohort of early-stage PD compared to HCs, using a flow cytometry panel designed to characterise peripheral immune changes across a broad range of innate and adaptive immune cell populations(Greenland et al., 2025). FOXP3 is a master transcription factor required for Treg identity and function, while HELIOS is expressed by roughly two thirds of Tregs and has been implicated in the maintenance of the regulatory phenotype(Sebastian et al., 2016; Shevyrev & Tereshchenko, 2020).

The potential therapeutic relevance of Tregs in PD has attracted increasing interest over recent years. Consistent with a potential protective role of Tregs, animal studies have reported that increased Treg numbers or adoptive Treg transfer can improve motor outcomes and reduce dopaminergic neurotoxicity(Badr et al., 2022; Bennett & Piercey, 1999; Guevara- Salinas et al., 2024; S.-Y. Park et al., 2023; Wohler et al., 2009; Xie et al., 2024; Yamamoto et al., 2022).

This has led to early clinical studies of approaches to expand Tregs in individuals with PD. One phase I clinical trial, led by Howard Gendelman, investigated sargramostim, a granulocyte-macrophage colony-stimulating factor (GM-CSF), as a means of modulating neuroinflammation by enhancing Treg numbers in people with PD. Treatment was associated with increased Treg numbers which correlated with improvements in MDS-UPDRS motor examination (part III) scores(Olson et al., 2021, 2023; Sil et al., 2026). A phase II study is planned.

In 2024, Coya Therapeutics announced a licence to develop low-dose IL-2 combined with GM-CSF to expand Tregs for the treatment of neurodegenerative diseases, including PD. Clinical trials of this combination in PD have not yet begun. However pre-clinical studies (also by Gendelman and colleagues) have shown that IL-2 mediated Treg expansion is protective in a mouse model of PD. The expanded Tregs expressed higher levels of suppressive markers, ICOS and CD39, and their adoptive transfer into MPTP-treated mice led to increased dopaminergic neuron growth in the substantia nigra and striatum. Data shared by Coya Therapeutics from the Gendelman lab also demonstrated that combining IL-2 and GM-CSF increased Treg numbers more than either intervention alone(*Coya Therapeutics Licenses Intellectual Property Rights for Use of Next Generation Immune Modulatory Biologics in Combination with COYA 301 to Enhance Regulatory T Cells (Tregs) in Inflammatory Diseases*, 2024.; Markovic et al., 2022). Coya Therapeutics are also developing other Treg-boosting therapies utilising IL-2 with CTLA-4-Ig fusion protein or GLP-1 receptor agonist for the treatment of neurodegenerative diseases(Read, 2024; Thome et al., 2026).

Evidence from other human studies is also emerging. A 2024 observational study of early-stage PD patients reported that treatment with pramipexole or levodopa was associated with an increase in circulating Tregs at one year, which correlated with improvements in Hoehn & Yahr and MDS-UPDRS scores(Arce-Sillas et al., 2024). Treg-based approaches may also have relevance to emerging cell-replacement therapies, with autologous Treg co-transplantation reported to protect dopaminergic grafts in a recent preclinical study(T.-Y. Park et al., 2023).

Despite growing interest in Tregs in PD, their phenotype and functional capacity remain unclear, with previous studies differing in their definitions of Tregs and assessment of suppressive function. Many previous studies have also failed to account for the effects of biological sex or levodopa treatment on Treg numbers and function(Álvarez-Luquín et al., 2019; Yang et al., 2016; Y. Chen et al., 2015). Understanding whether Tregs are altered in number or phenotype and whether they retain their suppressive function is of critical importance to therapeutic approaches aimed at expanding these cells, as their potential benefit will depend on the regulatory competence of the cells being expanded. We therefore used high-parameter spectral flow cytometry, epigenetic assessment of the Treg-specific demethylation region (TSDR) and direct functional assessment to determine whether Tregs are quantitatively or functionally altered in PD and to comprehensively characterise the cells that might be targeted by Treg-based therapies in much greater depth than previous studies.

## 2. Methods

### 2.1. Participants and sample collection

PD and HC participants were recruited through the Cambridge Parkinson’s Research Clinic. Ethical approval was obtained from the East of England – Cambridge Central Research Ethics Committee (Genetic, Metabolic and Immune Abnormalities in Neurodegenerative Disorders of the Central Nervous System, REC ref 03/303). Participants were excluded if they were receiving immunomodulatory therapy, had an autoimmune disease, had received a vaccination within the preceding 3 weeks, had taken ibuprofen within 48 hours of blood collection, or had a current infection or symptoms of a cold or influenza. Following informed consent, five 9ml lithium heparinised blood tubes were collected from each participant and processed within a few hours.

### 2.2. Peripheral blood mononuclear cell isolation

Blood samples were diluted 1:1 with PBS and layered onto Lymphoprep density gradient medium in SepMate tubes (both StemCell Technologies) followed by centrifugation at 1200g and room temperature for 10 minutes. The PBMC layer was collected and washed with PBS, centrifuging at 400g for 10 minutes three times, prior to downstream analysis. For phenotypic analysis, PBMCs were cryopreserved in 90% heat-inactivated foetal calf serum (FCS) and 10% dimethyl sulfoxide (DMSO), initially stored at -70°C in an ultra-low temperature freezer (ULTF) and transferred to liquid nitrogen within 1-3 days. Freshly isolated PBMCs were used for functional assays.

### 2.3. Spectral flow cytometric analysis for phenotyping

Cryopreserved PBMCs were thawed and stained with a 24-colour panel adapted from (Georgieva et al., 2024). This involved staining with viability dye for 15 minutes in the dark at room temperature first, followed by chemokine receptor staining under the same conditions. Next, the samples were stained on ice with antibodies for surface proteins in a mastermix of FACS buffer (100µl per sample) and incubated at 4°C for 45 minutes in the dark. The panel comprised: 1µl CD45RA-BUV395 (clone: HI100; BD Biosciences), 1:1500 LIVE/DEAD Blue (Invitrogen), 2µl CD3-BUV496 (clone: UCHT1; BD Biosciences), 2µl CD56-BUV563 (clone: NCAM16.2; BD Biosciences), 1µl CD39-BUV661 (clone: TU66; BD Biosciences), 1.25µl CD69-BUV737 (clone: FN50; BD Biosciences), 2µl CD73-BUV805 (clone: AD2; BD Biosciences), 1µl of each clone CD25-BV421(2 clones: MA251, 2A3; BD Biosciences), 0.625µl Ki67-eFluor450 (clone: 20Raj1; eBioscience/Thermo Fisher Scientific), 1.25µl CD4-V500 (clone: RPA-T4; BD Biosciences), 1µl CCR4-BV605 (clone: L291H4; Biolegend), 2.5µl CCR6-BV650 (clone: G034E3; Biolegend), 2.5µl PD-1-BV785 (clone: EH12.2H7; Biolegend), 5µl CXCR4-AF488 (clone: HIT2; R&D Systems), 1.25µl CD14-cFluorB548 (clone: 63D3; Cytek), 2µl CD8-PerCP (clone: SK1; Biolegend), 2µl CD19-BB700 (clone: SJ25C1; BD Biosciences), 1µl of each clone FOXP3-PE (2 clones: PCH101 and 259D; eBioscience/Thermo Fisher Scientific and Biolegend), 1µl HELIOS-PEDazzle (clone: 22F6; Biolegend), 14µl CTLA-4-PECy5 (clone: BNI3; BD Biosciences), 1µl CXCR3-PECy7 (clone: G025H7; Biolegend), 2µl CD27-APC (clone: MT271; Biolegend), 2µl CD127-APCR700 (clone: HIL-7R-M21; BD Biosciences), 5µl LFA-1-APC Fire750 (clone: m24; Biolegend).

BD Horizon Brilliant Stain buffer was used to minimise dye interactions during staining. Cells were fixed and permeabilised using the FOXP3/Transcription Factor Staining Buffer Set (eBioscience/Thermo Fisher Scientific) for 25 minutes at 4°C in the dark before staining with intracellular antibodies made in a mastermix with permeabilisation buffer (100µl per sample). Cells were incubated overnight with intracellular antibodies and washed the following morning, ready for analysis on the Cytek Aurora spectral cytometer.

Flow cytometry data were analysed using FlowJo v10. Lymphocytes were identified by forward-and side-scatter characteristics, followed by exclusion of doublets, dead cells, monocytes (CD14+), B cells (CD19+) and CD56+ NK/NKT cells. CD3 T cells were subsequently divided into CD4 and CD8 and further classified according to differentiation state. Naive cells were defined as CD27+CD45RA+, while memory cells comprised central memory (CM; CD27+CD45RA-), effector memory (EM; CD27-CD45RA-) and terminally differentiated effector memory cells re-expressing CD45RA (TEMRA; CD27-CD45RA+).

Tregs were defined as CD127lowFOXP3+ and T effectors as FOXP3-. Naive and memory Tregs were further characterised according to HELIOS and CD25 expression. Although CD25 is often used as a surface marker to define Tregs, CD25 low/negative FOXP3+ Tregs have been described in chronic inflammatory conditions(Ju et al., 2022; Quiroga et al., 2012). Additional markers (CCR4, CCR6, CD39, CD69, CD73, CXCR3, CXCR4, Ki67, LFA-1, CTLA-4 and PD-1) were assessed to further characterise the activation, differentiation and functional phenotype of Tregs and conventional T cells. The gating strategy is shown in **supplementary figure S1**.

### 2.4. Treg suppression assay

PBMCs were enriched for T cells using the PanT human cell isolation kit (Miltenyi Biotec), as per the manufacturer’s instructions. Enriched T cells were stained using Realease antibodies against: 2µl CD4-VioBlue (clone: REAL103), 2µl CD8-VioGreen (clone: REAL100), 2µl CD3-PerCP (clone: REAL104), 2µl CD127-PE (clone: REAL102), 2µl CD25-APC (clone REAL128) (Miltenyi Biotec), 1:400 Zombie NIR (Biolegend) (**supplementary figure S2**). The samples rested in FACS buffer at 4°C overnight and CD4 Tregs (CD4+CD127lowCD25high) and T effectors (CD3+CD25low) were subsequently isolated by fluorescence activated cell sorting (FACS) on a BD FACSAria Fusion.

Realease antibodies were released as per the manufacturer’s instructions (Miltenyi Biotec) and sorted cells rested in the incubator at 37°C for 1 hour in assay media (X-VIVO supplemented with 3% human AB serum and 1% penicillin-streptomycin). Tregs were stained with cell proliferation dye efluor670 and T effectors with efluor450 (eBioscience/Thermo Fisher Scientific) and then 50,000 T effectors were cultured with Tregs at ratios ranging from 1:1 to 16:1 (T effectors:Tregs) in 96 well V-bottom plates. T cell activation was induced using Treg suppression inspector beads (Miltenyi Biotec), according to the manufacturer’s instructions, assay media.

Following 5 days in culture at 37°C, cells were stained with the following antibodies: 1µl CD3-BUV395 (clone: UCHT1; BD Biosciences), 1µl CD4-FITC (clone: OKT4; Biolegend), 1µl CD8-PECy7 (clone: SK1; Biolegend), 1:400 Zombie red (Biolegend) (**supplementary figure S3**) and analysed on a BD Fortessa 2. Percentage suppression was calculated using the equation:

(Proliferation of T effectors alone — proliferation of T effectors with Tregs)/ proliferation of T effectors alone x100

Technical replicates were included where sufficient cell numbers permitted.

### 2.5. Cytokine Analysis

Cytokines were measured in supernatants from the Treg suppression assay using the Bio-Plex Pro Human Cytokine 8-plex plate (Bio-Rad) with the addition of IL-1β and IL-17A (Bio-Rad). Samples were diluted 1:2 in assay media and the assay performed according to manufacturer’s instructions. Cytokine suppression was calculated using the equation: (Concentration for T effectors alone — concentration for T effectors with Tregs)/concentration for T effectors alone x100

To further investigate IL-17A production, CD4 and CD8 T effectors were isolated by FACS using the sorting strategy described above and stimulated at 50,000 cells per well with anti-CD3/CD28 Dynabeads (concentration of 1µl per 1.2 million T cells) (Thermo Fisher Scientific) for 1, 3 or 5 days. IL-17A concentrations in culture supernatants were measured by ELISA (Human IL-17A [homodimer] uncoated ELISA kit; Thermo Fisher Scientific), according to the manufacturer’s instructions. Cells were collected for gene expression analysis and stored in RNA lysis buffer (Zymo Research) at -70°C.

RNA was extracted using the Quick-RNA Microprep Kit (Zymo Research) following DNAse treatment and reverse transcribed to cDNA using polyT primer. Expression of *IL-17A*, *IL-17F* and *RORC* was quantified by quantitative PCR (qPCR) using primers designed spanning exon-exon junctions: *IL-17A*_forward AACCGATCCACCTCACCTTG, *IL-17A*_reverse GCACTTTGCCTCCCAGATCA, *IL-17F*_forward GAAAACCAGCGCGTTTCCAT, *IL-17F*_reverse GCAGCCCAAGTTCCTACACT, *RORC*_forward AGCGCTCCAACATCTTCTCC, *RORC*_reverse CACCACGTACTGAATGGCCT. *β2-microglobulin* was used as the reference gene and primers purchased from Thermo Fisher Scientific, B2M qPCR assay (Hs00187842_m1). Samples were analysed in duplicate.

### 2.6. Treg-specific demethylation region (TSDR) analysis

PBMCs from 6 participants with PD and 6 HC participants underwent T cell enrichment and were stained using the following antibodies for FACS sorting: CD3 BUV395 (BD Biosciences), CD4-BV650 (BD Biosciences), CD8-V500 (BD Biosciences), CD25-BV421 (BD Biosciences), CD127-PECy7 (Biolegend), CD45RA-BV785 (Biolegend), CD27-APC (Biolegend) and Zombie NIR (Biolegend). Four populations were sorted: CD4 naive Tregs (CD4+CD45RA+CD27+CD127lowCD25high), CD4 naive T effectors (CD4+CD45RA+CD27+CD25low), CD4 memory Tregs (comprising CM; CD4+CD45RA-CD27+CD127lowCD25high, EM; CD4+CD45RA-CD27-CD127lowCD25high and TEMRA; CD4+CD45RA+CD27-CD127lowCD25high) and CD4 memory T effectors (comprising CM; CD4+CD45RA-CD27+CD25low, EM; CD4+CD45RA-CD27-CD25low, and TEMRA; CD4+CD45RA+CD27-CD25low) (**supplementary figure S4**).

Treg-specific demethylation region (TSDR) methylation was assessed using the protocol described by Rainbow and colleagues (Rainbow et al., 2015), with sequencing performed at the CRUK Cambridge Institute. As the TSDR is located on the X chromosome, values from female participants were adjusted to account for X-inactivation by doubling the measured demethylation percentage, allowing comparison of TSDR demethylation between male and female participants.

### 2.7. Statistical analysis

Statistical analyses were performed using R (version 2024.12.0+467) and GraphPad Prism (version 10.6.0). Data distribution was assessed using the Shapiro-Wilk test, and statistical tests were selected accordingly. For comparisons between two groups, parametric or non-parametric tests were used as appropriate. Statistical significance was defined as p<0.05. Graphs denote significance as follows: *=p<0.05, **=p<0.01.

Given the exploratory nature and high dimensionality of the spectral flow cytometry analysis, individual phenotypic markers were analysed without adjustment for multiple comparisons. These analyses were interpreted in the context of the overall phenotypic pattern rather than as independent hypothesis tests.

## 3. Results

### 3.1. Participant demographics

In total, 29 participants with PD and 26 healthy controls were recruited. Participant demographics and numbers included in each assay are summarised in the tables below.

**Table 1.**
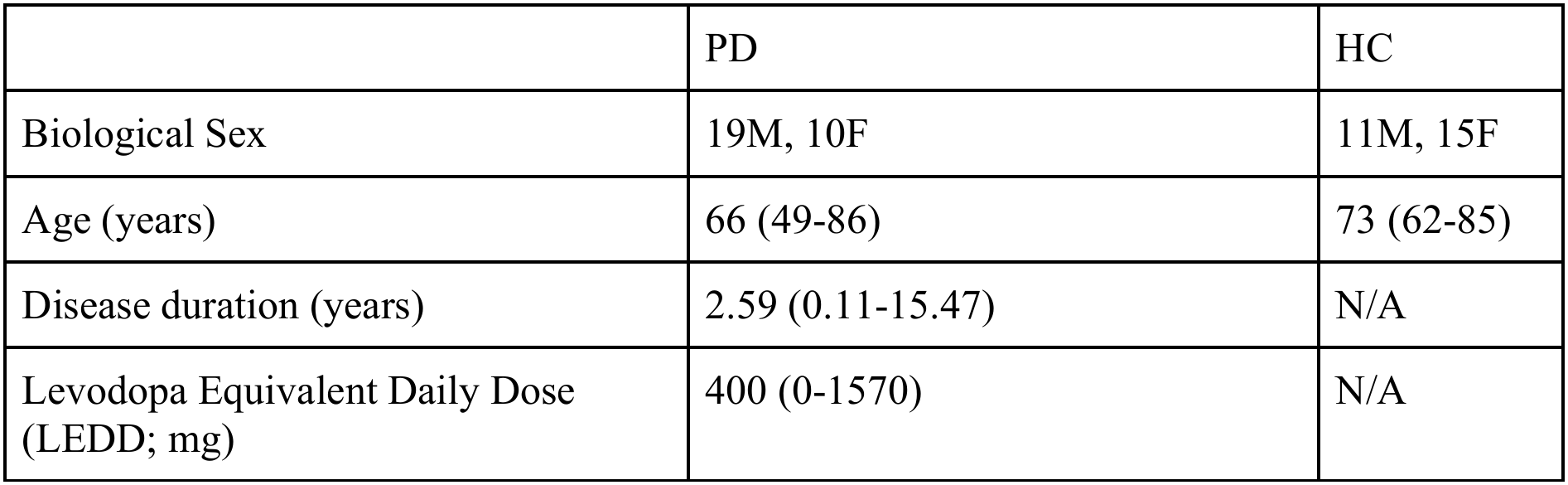
Demographics of participants phenotyped by spectral flow cytometry. Data shown as median with range in brackets.

|  | PD | HC |
|---|---|---|
| Biological Sex | 19M, 10F | 11M, 15F |
| Age (years) | 66 (49-86) | 73 (62-85) |
| Disease duration (years) | 2.59 (0.11-15.47) | N/A |
| Levodopa Equivalent Daily Dose (LEDD; mg) | 400 (0-1570) | N/A |

**Table 2.** Demographics of participants for functional analysis using suppression assays. Data shown as median with range in brackets.

|  | PD | HC |
|---|---|---|
| Biological Sex | 5M, 0F | 1M, 4F |
| Age (years) | 78 (64-82) | 73 (63-78) |
| Disease duration (years) | 3.74 (0.18-15.47) | N/A |
| LEDD (mg) | 300 (160-1570) | N/A |

### 3.2. T effector and Treg proportions are similar between PD and HC

Spectral flow analysis of cryopreserved PBMCs showed no significant differences in the proportions of total CD3 T cells, CD4 and CD8 T cells between PD and HC. Within both the CD4 and CD8 compartments, the proportions of naive and memory cells were also similar between groups. Similarly, no significant differences were observed in the proportions of naive or memory T effectors or Tregs; CD4 naive Tregs 1.52%(±0.68) PD vs 2.15%(±1.66) HC, CD4 memory Tregs 5.50%(±2.06) PD vs 5.83%(±2.29) HC, CD8 naive Tregs 0.06% (±0.05) PD vs 0.09%(±0.14) HC, CD8 memory Tregs 0.41%(±0.33) PD vs 0.40%(±0.41) HC (**figure 1**).

**Figure 1.**
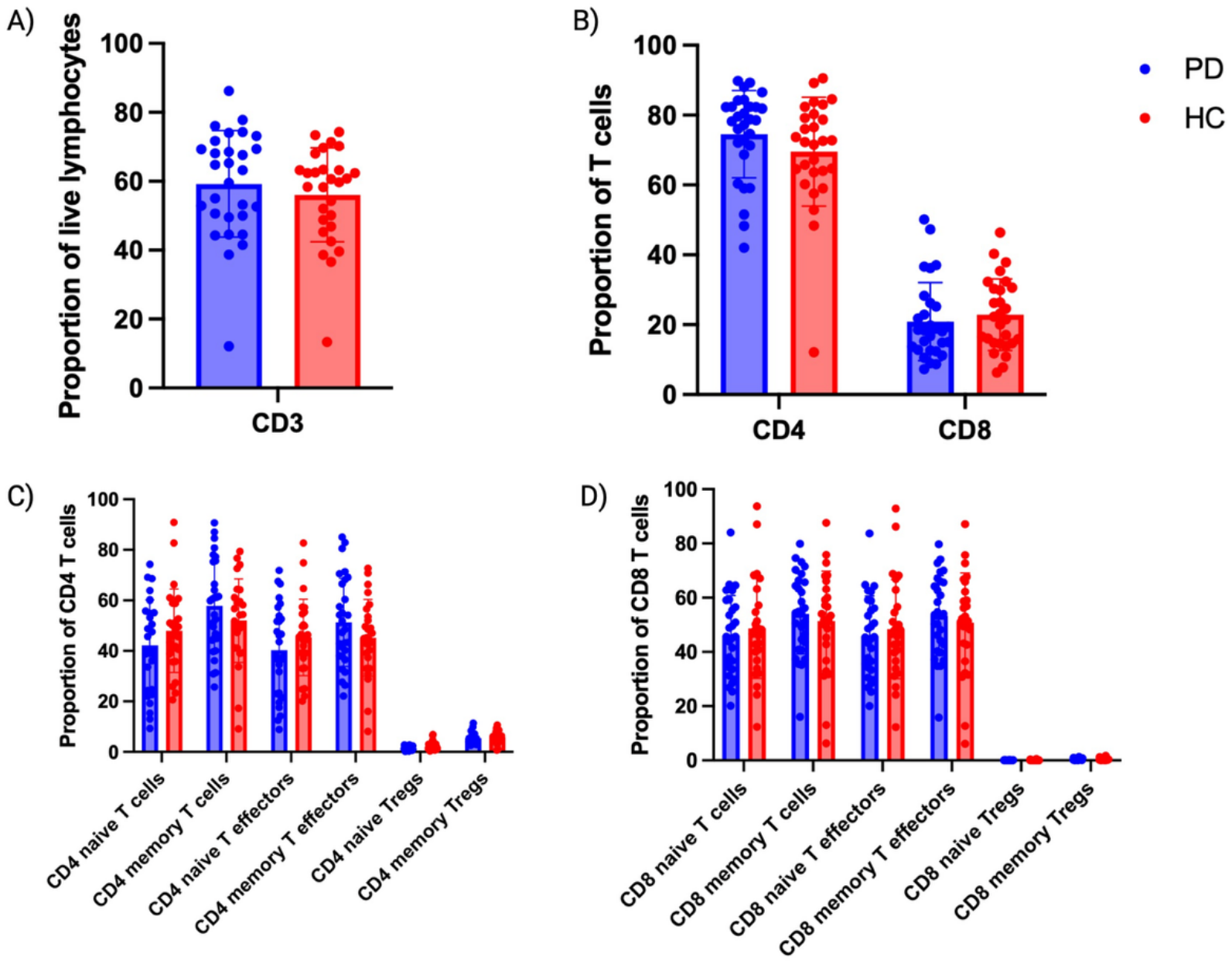
Proportions of T cells and CD4 subsets. **A)** CD3 T cells as a proportion of live lymphocytes. **B)** CD4 and CD8 T cells as a proportion of total T cells. **C)** CD4 naive and memory populations as a percentage of CD4 T cells. **D)** CD8 naive and memory populations as a percentage of CD8 T cells. Graphs show mean ±SD. PD, n = 29; HC, n = 26.

Since PD is a male-dominated disease and given potential sex-related differences in immune phenotype, the proportion of naive and memory Tregs was compared between sex-matched PD and HC groups. Tregs were defined as CD127lowFOXP3+ and either naive (CD27+CD45RA+) or memory (through a not-naive gate). No significant differences were observed in either Treg population in males or females (**supplementary figure S5**).

The proportion of naive or memory Tregs was also not associated with disease duration or levodopa-equivalent daily dose (LEDD). Consistent with these findings, mixed-effects modelling incorporating both disease duration and LEDD identified no significant association of either variable with the proportion of naive or memory Tregs. (**supplementary figure S6**).

### 3.3. CD4 memory Tregs are similar between PD and HC

Having found no quantitative differences in circulating CD4 Treg populations, including after consideration of sex, disease duration and treatment exposure, we next examined their phenotype in greater detail.

The proportions of CD4 memory Tregs expressing HELIOS and CD25, including HELIOS+CD25+, HELIOS+CD25-, HELIOS-CD25+ and HELIOS-CD25-populations did not significantly differ between PD and HC. Similarly, the expression levels of FOXP3, HELIOS and CD25, assessed by mean fluorescence intensity (MFI) among marker-positive cells, were comparable between groups (**figure 2A-E**).

**Figure 2.**
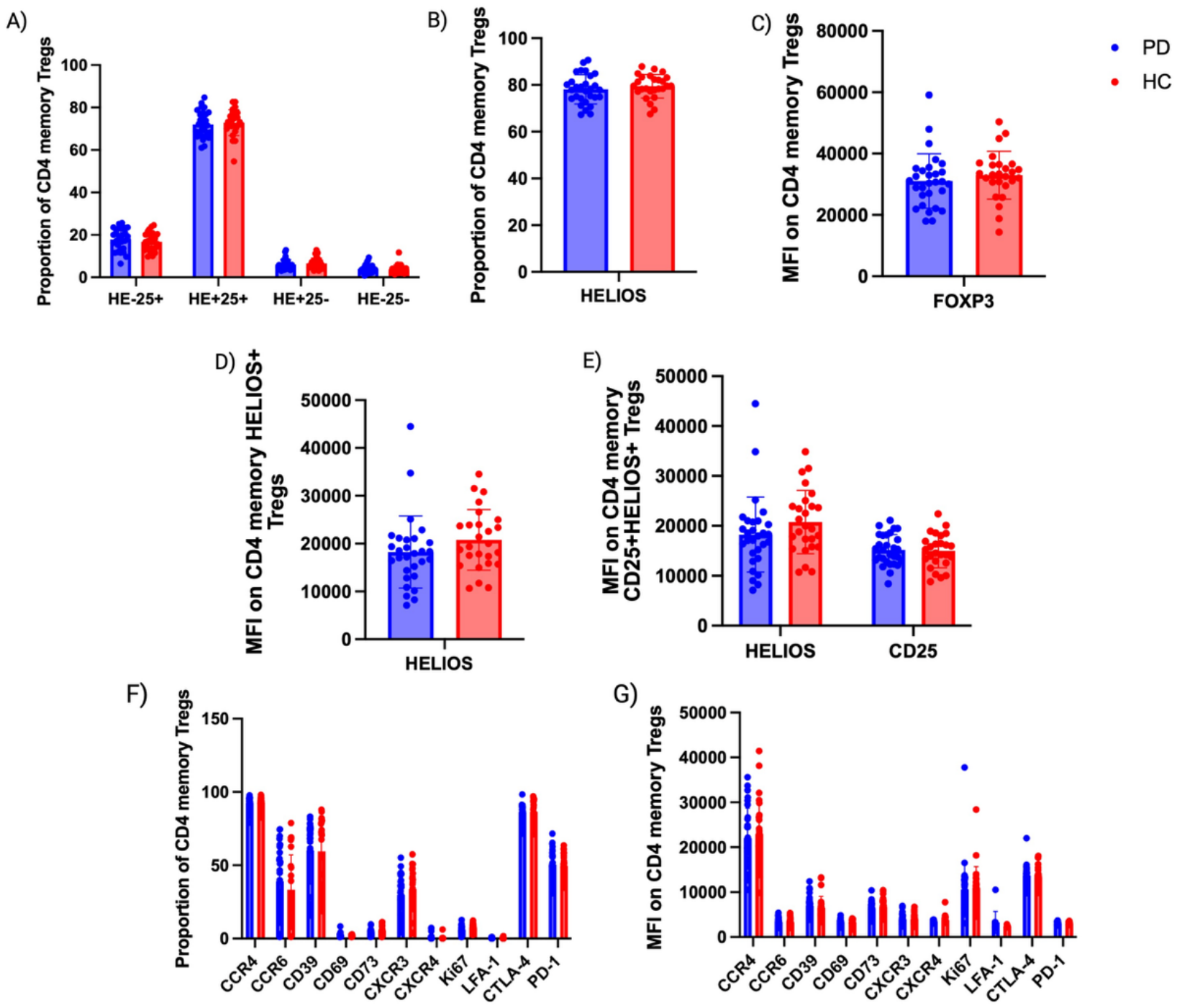
Expression of phenotyping markers on CD4+CD127lowFOXP3+ memory Tregs. **A)** Proportion of subpopulations within the CD127lowFOXP3+ memory Treg pool, defined by HELIOS (HE) and CD25 (25) expression. **B)** Total HELIOS expression as a proportion of CD4 memory Tregs. **C)** FOXP3 MFI of CD4 memory Tregs. **D)** HELIOS MFI of CD4 memory HELIOS+ Tregs. **E)** HELIOS and CD25 MFI of CD25+HELIOS+ CD4 memory Tregs. **F)** Proportions of CD4 memory Tregs expression downstream Treg markers **G)** MFI of downstream Treg markers in marker-positive CD4 memory Tregs. Graphs show mean ±SD. PD, n = 29; HC, n = 26.

We next assessed 11 additional markers associated with Treg activation, differentiation and functional phenotype: CCR4, CCR6, CD39, CD73, CD69, CXCR3, CXCR4, Ki67, LFA-1, CTLA-4, PD-1. No significant differences were observed between PD and HC in either the proportion of CD4 memory Tregs expressing these markers or their expression levels (MFI) among marker-positive cells. Overall, CD4 memory Tregs in PD showed a broadly similar phenotypic profile to those in HC, with no evidence of altered expression of the canonical Treg, activation or functional markers examined (**figure 2F-G**).

### 3.4. Reduction in HELIOS expression in naive CD4 Tregs in PD

Next, we examined CD4 naive Tregs in greater detail. The relative proportions of HELIOS+CD25+, HELIOS+CD25-, HELIOS-CD25+, HELIOS-CD25-, as well as the proportion of HELIOS+ CD4 naive Tregs within the total naive CD4+FOXP3+ Treg pool, did not differ significantly between PD and HC (**figure 3A-B**).

**Figure 3.**
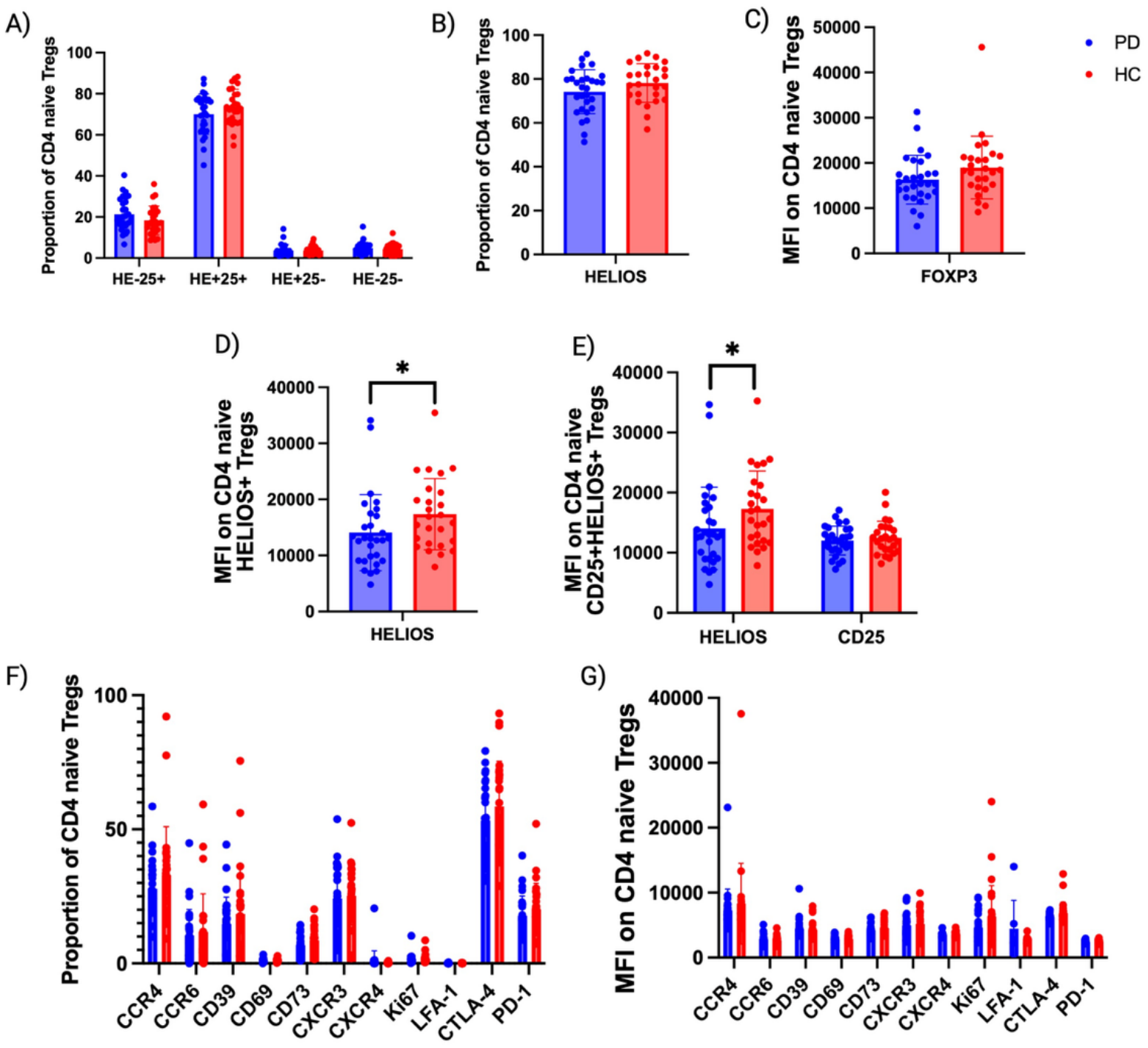
Expression of phenotyping markers on CD4+CD127lowFOXP3+CD27+CD45RA+ naive Tregs. **A)** HELIOS and CD25 expression as a proportion of CD4 naive Tregs. **B)** HELIOS expression as a proportion of CD4 naive Tregs. **C)** FOXP3 MFI of CD4 naive Tregs. **D)** HELIOS MFI of CD4 naive HELIOS+ Tregs. **E)** HELIOS and CD25 MFI of CD25+HELIOS+ CD4 naive Tregs. **F)** Proportions of CD4 naive Tregs expressing downstream Treg markers. **G)** MFI of downstream Treg markers in marker-positive CD4 naive Tregs. *p<0.05. Graphs show mean ±SD. PD, n = 29; HC, n = 26.

However, analysis of marker expression intensity revealed reduced HELIOS expression in PD. HELIOS MFI was reduced by 1.23-fold in both total HELIOS+ CD4 naive Tregs and the HELIOS+CD25+ subset compared with HC (p=0.0330 and p=0.03510, respectively). FOXP3 MFI of all CD4 naive Tregs showed a similar trend towards lower expression in PD (1.17-fold decrease), although this did not reach statistical significance (p=0.0781) (**figure 3C-E**). This result was sensitive to individual observations: exclusion of three identified outliers (2 PD, 1 HC), defined as values ±2 standard deviations from the mean, resulted in a statistically significant difference (p=0.0303).

To assess whether differences in sex distribution between the cohorts contributed to these findings, FOXP3 and HELIOS MFI were compared separately in male and female participants. No significant differences between PD and HC were observed within either sex, providing no evidence that the overall findings were attributable to sex (**supplementary figure S7**).

Analysis of the additional Treg-associated markers revealed no significant differences between PD and HC in either the proportion of marker-positive cells or MFI for CCR4, CCR6, CD39, CD69, CD73, CXCR3, CXCR4, Ki67, LFA-1, CTLA-4 or PD-1 (**figure 3F-G**). Thus, despite similar frequencies and subset distribution, naive CD4 Tregs in PD showed a selective reduction in HELIOS expression, with a similar but non-significant trend for FOXP3, rather than a broad alteration in their phenotypic profile.

### 3.5. Reduction in HELIOS MFI in naive CD8 Tregs in PD

The proportions of naive and memory CD8 T effectors and Tregs did not differ significantly between PD and HC. Similarly, there were no significant differences in the proportions of naive or memory CD8 Tregs expressing HELIOS and CD25, including HELIOS+CD25+, HELIOS+CD25-, HELIOS-CD25+, HELIOS-CD25-, or in the overall proportion of HELIOS+ CD8 Tregs (**figure 4A-D**).

**Figure 4.**
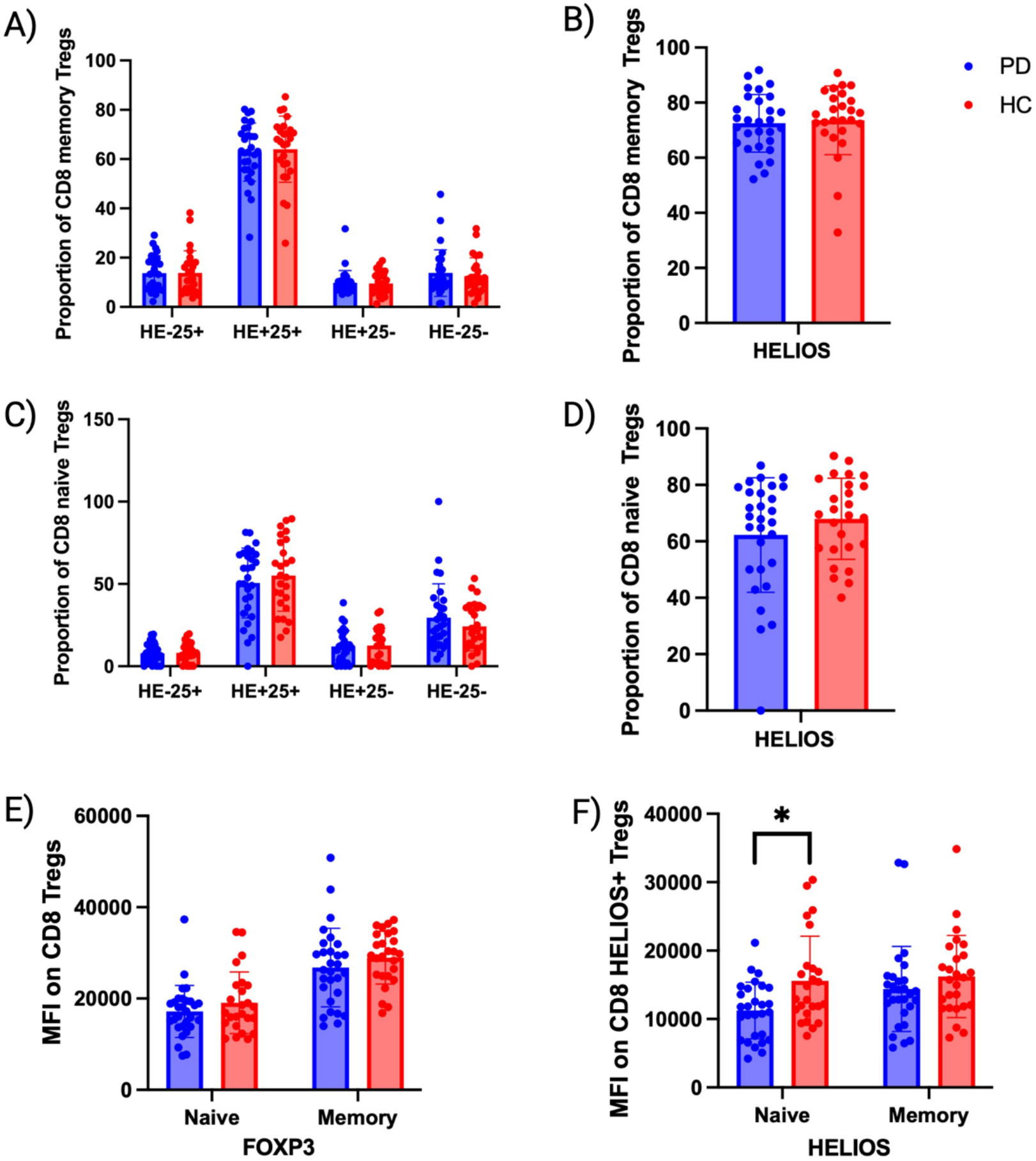
CD8 Tregs. **A)** HELIOS and CD25 expression as a proportion of CD8 memory Tregs. **B)** HELIOS+ CD8 memory Tregs as a proportion of CD8 memory Tregs. **C)** HELIOS and CD25 expression as proportions of CD8 naive Tregs. **D)** HELIOS+ CD8 naive Tregs as a proportion of CD8 naive Tregs. **E)** FOXP3 MFI on CD8 naive and memory Tregs. **F)** HELIOS MFI on HELIOS+ CD8 memory and naive Tregs. *p<0.05. Graphs show mean ±SD. PD, n = 29; HC, n = 26.

Given the differences in HELIOS expression observed in the CD4 Treg compartment, HELIOS and FOXP3 expression levels were also assessed by MFI in CD8 Tregs. No significant differences were observed in FOXP3 MFI in naive or memory CD8 Tregs, or in HELIOS MFI in memory CD8 Tregs. However, HELIOS MFI was significantly reduced by 1.39-fold in CD8 naive Tregs in PD compared to control (p=0.0133)(**figure 4E-F**). This finding should be interpreted cautiously, as naive CD8 Tregs are rare, with a mean of only 120 cells captured per donor.

### 3.6. CD4 Tregs show similar FOXP3 TSDR demethylation in PD and HC

Treg-specific demethylation region (TSDR) analysis showed no significant differences in *FOXP3* TSDR demethylation between PD and HC in either naive or memory CD4 Tregs. There were also no significant differences in TSDR demethylation between naive and memory Tregs within either the PD or HC groups (**figure 5A-B**). These findings indicate that the epigenetic status of the *FOXP3* locus, and thus Treg phenotypic stability, was comparable in CD4 Tregs from PD and HC. As a method to validate the purity of the naive Treg pool, the proportion of surface gated naive Tregs (CD127lowCD25high) that were FOXP3+ was calculated from the spectral data. This gave a mean of 91.80% (median 92.63%) (**figure 5C**). Since the sort gates were conservative, ensuring only the brightest CD25+ Tregs were selected, this calculation gave fair assurance that the naive Treg population sorted was a pure representative of Tregs with very few contaminating T effectors.

**Figure 5.**
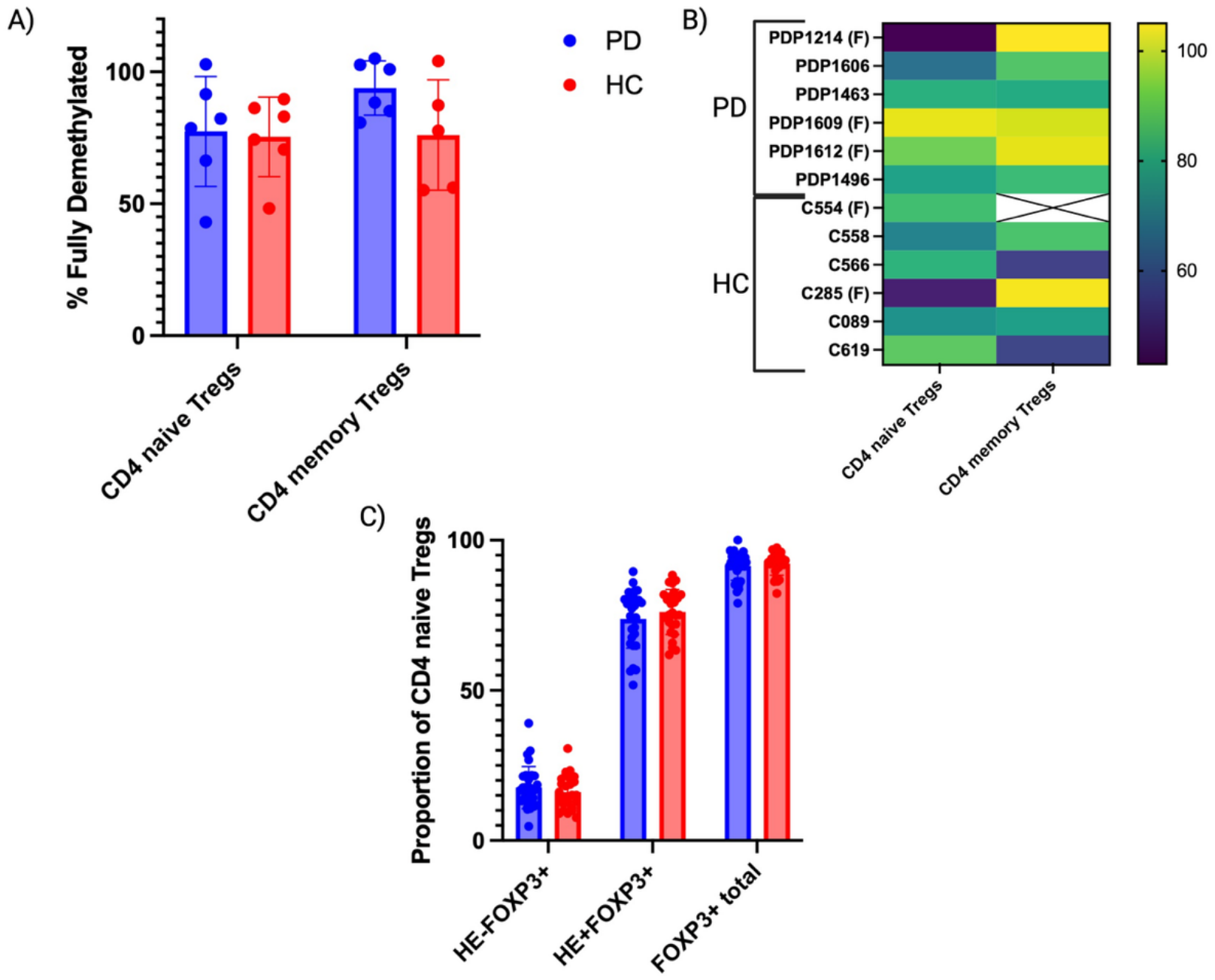
FOXP3 demethylation status of naive and memory CD4 Tregs. **A)** Comparison of TSDR demethylation in naive and memory Tregs between PD and HC groups. **B)** Heatmap of all donors showing the percentage of fully demethylated cells for each donor. Donor C554 had no CD4 memory Tregs sample. Female donors are marked with an (F). **C)** Proportion of surface gated CD4 naive CD127lowCD25high Tregs expressing FOXP3 including HELIOS+ and HELIOS-. Graphs show mean ±SD. PD, n = 6; HC, n = 6.

### 3.7. CD4 Tregs show similar suppressive function in PD and HC

The suppressive capacity of CD4 Tregs was assessed using a polyclonally stimulated bead-based suppression assay with autologous T effectors. No significant differences in suppressive capacity were observed between PD and HC Tregs. Suppression of bulk CD3, CD4 and CD8 T effector proliferation was comparable between groups across the T effector ratios tested (**figure 6**). Thus, CD4 Tregs from PD demonstrated preserved suppressive function *in vitro* compared with HC Tregs.

**Figure 6.**
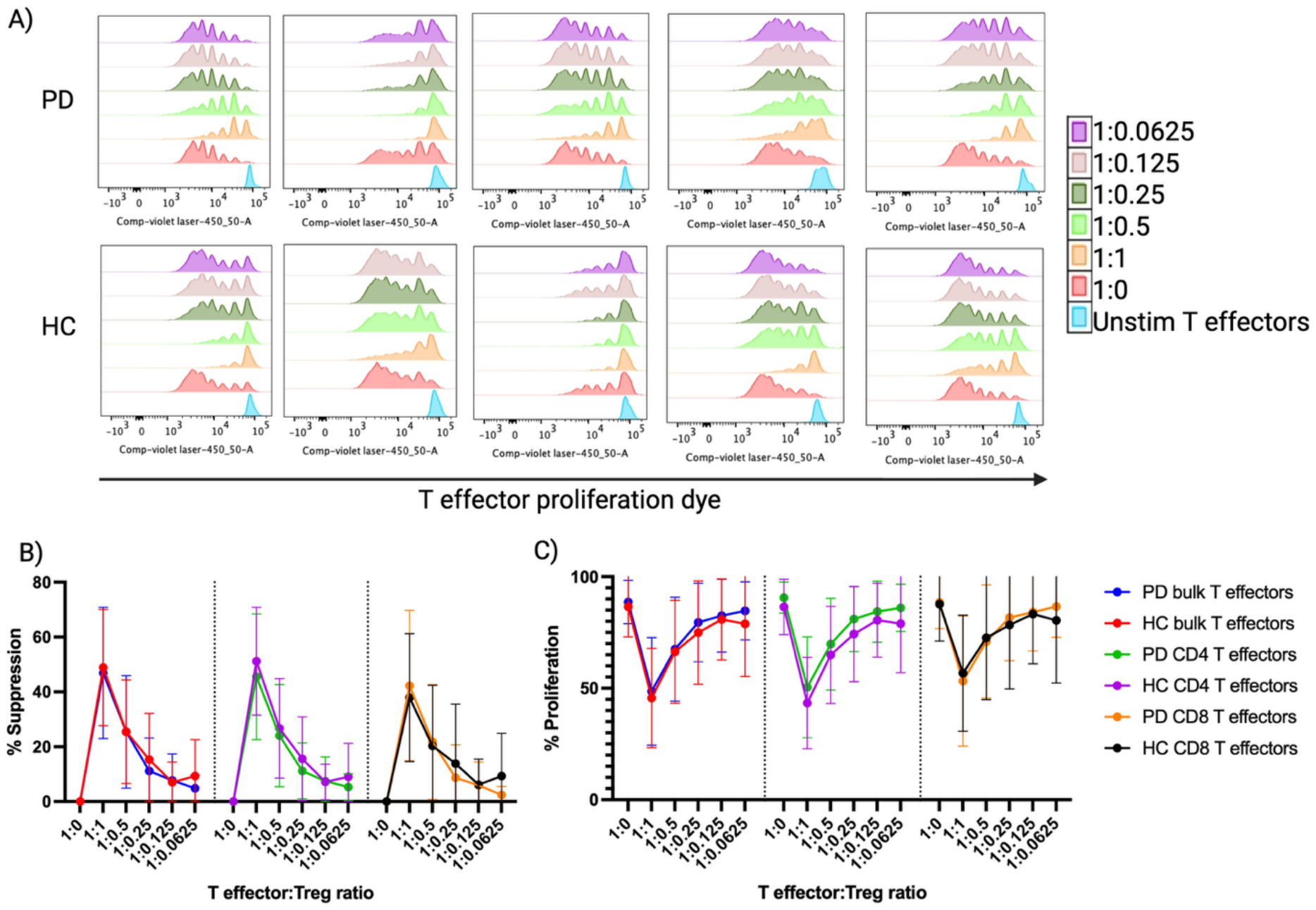
Autologous suppression assays. **A)** Proliferation histograms, showing CD3 T effectors from all donors proliferated and were suppressed by their autologous Tregs. **B)** Suppression of bulk, CD4 and CD8 autologous T effectors at different ratios of T effectors:Tregs. **C)** Proliferation of bulk, CD4 and CD8 autologous T effectors at different ratios of T effectors:Tregs. Technical replicates were averaged for each donor and timepoints. Graphs show mean ±SD. PD, n = 5; HC, n = 5.

### 3.8. Cytokine suppression by Tregs and production by T effectors is broadly similar between PD and HC

To determine whether Tregs from PD and HC differentially suppressed distinct cytokine responses, supernatants from the Treg suppression assays were analysed for cytokines associated with Th1 (IL-2, TNF-α, IFN-γ, IL-8, IL-1β), Th2 (IL-4, IL-6) and Th17 (IL-17) responses, as well as the immunoregulatory/Th2 cytokine IL-10. No significant differences in cytokine proliferation or suppression were observed between PD and HC across the T effector ratios tested for any of the cytokines analysed (**figure 7A**). IL-1β was below the limit of detection for all samples.

**Figure 7.**
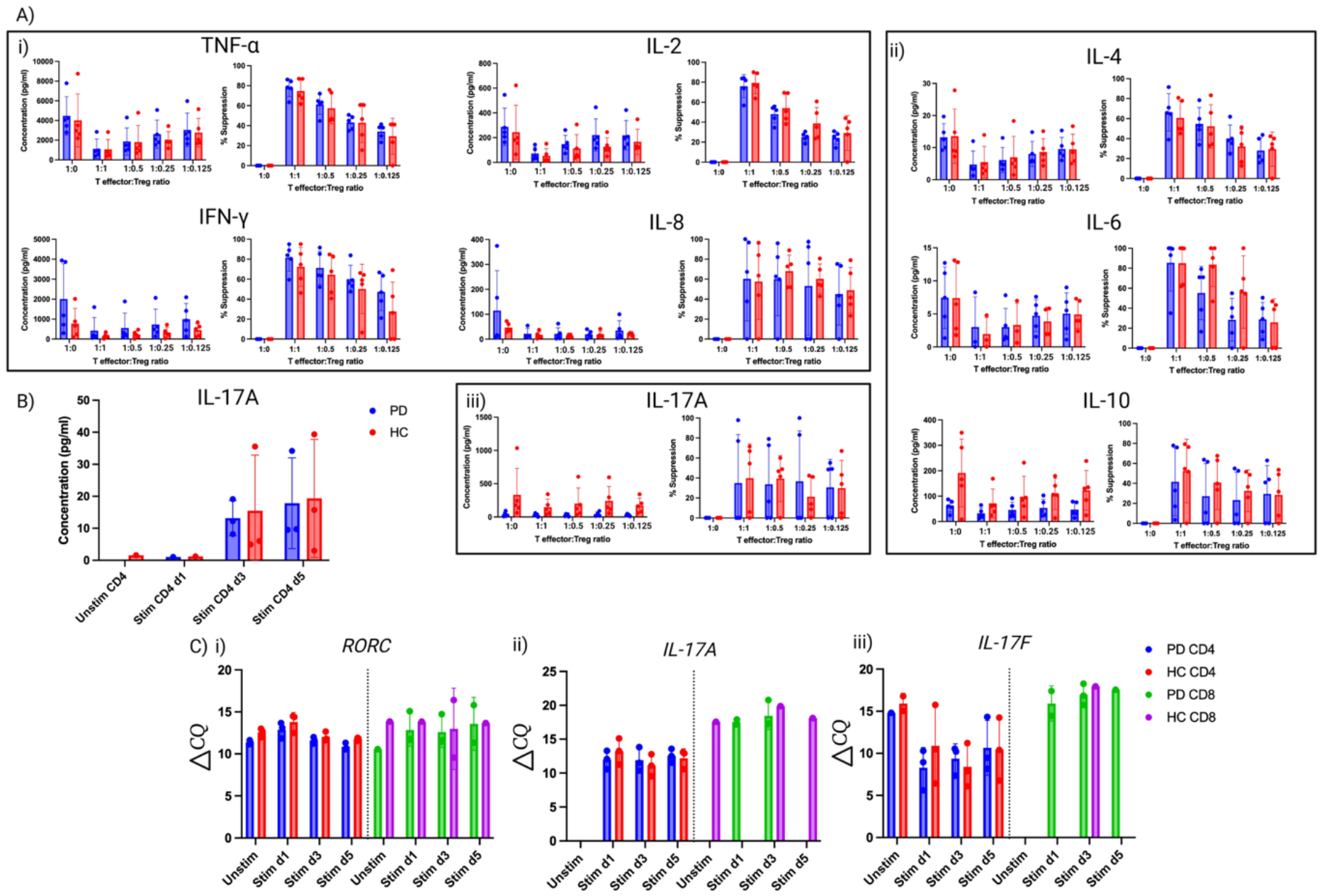
Cytokine analysis. **A)** Cytokine concentrations in suppression assay supernatants and their corresponding degree of suppression **i)** Th1 cytokines: TNF-α, IL-2, IFN-γ, IL-8. **ii)** Th2 cytokines: IL-4, IL-6, IL-10. **iii)** Th17 cytokine: IL-17A. PD, n = 5; HC, n = 5. **B)** IL-17A concentrations in stimulated CD4 T effector cultures, measured by ELISA. PD, n = 3; HC, n = 3. **C)** qPCR quantification of mRNA, normalised to B2M (ΔCQ), in T effectors from stimulation assay. **i)** RORC. **ii)** IL-17A. **iii)** IL-17F. Graphs shown mean ±SD. PD, n = 3; HC, n = 3.

We next examined cytokine concentrations in the T effector-only cultures (1:0), allowing cytokine production to be assessed in the absence of Treg-mediated suppression. PD T effectors showed trends towards higher concentrations of IFN-γ and IL-8 and lower concentrations of IL-17A and IL-10 compared with HC, although none of these differences reached statistical significance. Notably, IL-17A concentrations appeared particularly low in the PD group. Given this observation, CD4 and CD8 T effectors were subsequently stimulated independently for 1, 3 or 5 days to further investigate IL-17A production. IL-17A was not detectable by ELISA in any CD8 T effector supernatant. In CD4 T effectors, IL-17A was detectable following 3 and 5 days of stimulation in all donors, whereas detection was limited at baseline and after 1 day of stimulation (**figure 7B**). IL-17A concentrations did not differ significantly between PD and HC at either day 3 or day 5, nor between days 3 and 5 within either group.

Expression of *IL-17A*, *IL-17F* and *RORC* was also assessed by qPCR in CD4 T effectors (**figure 7C**). *IL-17A* and *IL-17F* expression did not differ significantly between PD and HC at any time point. *IL-17A* was undetectable in unstimulated CD4 T effectors. *RORC* expression was higher in PD than HC in unstimulated and day 5 stimulated CD4 T effectors, as indicated by lower ΔCq values; however, these differences did not remain significant following correction for multiple comparisons. CD8 T effectors showed a similar pattern for *RORC*, but donor numbers were insufficient for statistical comparison.

## 4. Discussion

This study found no significant differences between individuals with PD and HCs in the proportions of naive and memory CD4 T cells, CD4 T effectors or Tregs. Importantly, this was accompanied by broadly preserved Treg phenotype, *FOXP3* TSDR demethylation and suppressive function. Together, these findings provide no evidence for a major quantitative or functional deficit in circulating Tregs in PD. However, detailed phenotyping identified reduced HELIOS expression in naive CD4 Tregs, with a similar directional trend in FOXP3 expression. Thus, rather than a simple loss of Tregs or their suppressive capacity, PD may be associated with more subtle changes in Treg phenotype.

A strength of our study is the depth of phenotypic characterisation of circulating Tregs. Rather than relying on a limited phenotypic definition, we used high-parameter spectral flow cytometry to examine Treg differentiation state, HELIOS and CD25 expression, and a broad range of markers associated with activation, trafficking and regulatory function. This was complemented by assessment of *FOXP3* TSDR demethylation and direct measurement of suppressive function. In addition, the study population was carefully selected, with participants receiving immunomodulatory therapy or with relevant inflammatory conditions excluded, and the potential effects of sex, disease duration and dopaminergic treatment considered in the analyses.

The observed reduction in HELIOS expression is consistent with Greenland et al. (2025), who reported lower HELIOS MFI in CD4 HELIOS+ Tregs in PD compared with age-and sex-matched HCs. Their analysis did not distinguish between naive and memory Tregs, whereas our study identifies reduced HELIOS expression specifically in the naive CD4 Treg compartment. Greenland et al. also reported reduced FOXP3 MFI and HELIOS MFI in CD8

Tregs in PD, without differences in CD8 Treg proportions, broadly consistent with our findings. However, these observations should be interpreted cautiously in both studies given the low numbers of CD8 Tregs available for analysis. In our study, there were on average only 120 naive CD8 Tregs per sample, of which 91 were HELIOS+, compared with 548 memory CD8 Tregs, of which 396 were HELIOS+. Nevertheless, the finding of altered HELIOS expression across both CD4 and CD8 Tregs in two independent studies supports the view that altered HELIOS expression may be a feature of Tregs in PD.

The biological significance of reduced HELIOS expression in naive Tregs remains unclear. HELIOS is an important component of the Treg transcriptional programme and has been implicated in the maintenance of Treg identity and stability(Sebastian et al., 2016; Thornton et al., 2019). The potential relevance of the naive Treg compartment to disease is also supported by (Ohkura et al., 2020) who found that genetic variants associated with autoimmune disease susceptibility were enriched in Treg-specific demethylated regions and super-enhancers in naive Tregs. Although this study did not investigate PD, it highlights the potential importance of changes in naive Tregs in disease. The similar directional reduction in FOXP3 expression may also be relevant, although this did not reach statistical significance.

One possibility is that the inflammatory environment in PD influences Treg phenotype. IL-6 and IL-27 can modulate HELIOS expression and Treg suppressive capacity experimentally(Bin Dhuban et al., 2019), and IL-6 has been reported to be elevated in PD and associated with non-motor symptoms(Diaz et al., 2022; Fu et al., 2023). Evidence for altered IL-27 in PD is less consistent(Kouchaki et al., 2018; Wan et al., 2024). An alternative possibility is that altered HELIOS expression reflects increased Treg plasticity, with naive Tregs more readily adopting effector phenotypes under inflammatory conditions and potentially contributing to inflammation. These possibilities are not mutually exclusive but remain speculative as serum cytokines and Treg plasticity were not assessed in this study.

The phenotypic difference in HELIOS expression was not accompanied by evidence of impaired Treg function. *FOXP3* TSDR demethylation was similar in naive and memory Tregs between groups, and bulk (naive and memory) CD4 Treg suppression assays showed preserved suppression of T cell proliferation and cytokine production. However, because the suppression assays assessed both naive and memory Tregs combined, they cannot exclude a functional difference within the smaller naive Treg population in which reduced HELIOS expression was observed. Thus, our findings argue against a major functional deficit in circulating Tregs, but do not exclude more subtle differences in specific Treg subsets or changes that emerge under inflammatory conditions. Furthermore, circulating Tregs provide only a limited view of immune regulation in PD, as changes in activation, trafficking or tissue localisation may not be reflected in peripheral blood.

The analysis of CD4 T effector cells was exploratory, as the panel was primarily designed to investigate the Treg compartment. Although cytokine analysis suggested some differences in cytokine production by PD T effectors, these were not statistically significant, and the apparent difference in IL-17A production was not confirmed in subsequent analysis of stimulated CD4 T cells. Overall, there was no evidence from this dataset for a major shift in circulating CD4 T cell composition.

## 5. Conclusion

In conclusion, Tregs remain an attractive therapeutic target in PD, with preclinical and some early clinical studies reporting improved outcomes following interventions that increase Treg numbers. Our findings do not argue against this approach; indeed, the absence of a major circulating Treg deficit may be encouraging, suggesting that increasing Treg numbers could augment a population with preserved regulatory capacity. However, the reduced HELIOS expression we observed provides a note of caution, as expanded Tregs will need to retain a stable phenotype and suppressive function, particularly in inflammatory environments and within tissue. Thus, therapeutic benefit may depend not simply on increasing Treg numbers, but on ensuring that the expanded cells remain functionally competent where they are needed.

## Acknowledgements

We are very grateful to all the participants who took part in this study. This work was supported by a Cambridge School of Clinical Medicine’s Doctoral Training Programme in Medical Research Fellowship awarded to SC, the NIHR Biomedical Research Centre funding to JJ (BRC-1215-20014) and the NIHR Cambridge Biomedical Research Centre (NIHR203312). The views expressed are those of the author(s) and not necessarily those of the NIHR or the Department of Health and Social Care. We thank the NIHR Cambridge Biomedical Research Centre Cell Phenotyping Hub and the Cancer Research UK Cambridge Institute for their support. We would also like to thank Professor Roger Barker for allowing us to work under his existing ethical approval.

## Author contributions

SC wrote the manuscript and carried out all experiments and analysis. AP, LJ, ZG, and DR were involved in experiment design. CWG supplied all donor samples. JJ oversaw overall study design. All authors contributed to editing of the manuscript.

## Notes

### Competing Interest Statement

The authors have declared no competing interest.

