## Supplementary figures for "Circulating CD4 regulatory T cells are broadly preserved in Parkinson’s disease despite altered HELIOS expression"

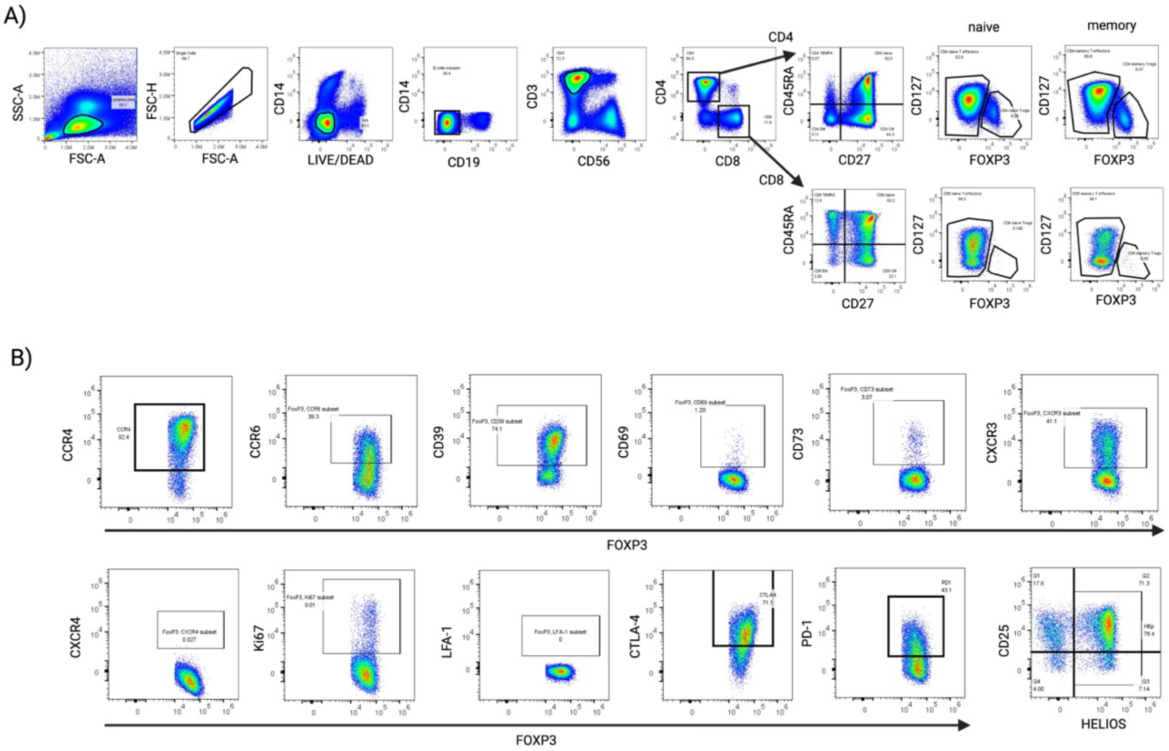


**S1.** ***Spectral flow phenotyping gating trees. A)*** *Sequential gating strategy used to identify memory and naive CD4 and CD8 Tregs and T effectors.* ***B)*** *Downstream marker gating, shown here for memory CD4 Tregs, with the same gating applied to naive Tregs and T effector populations.*

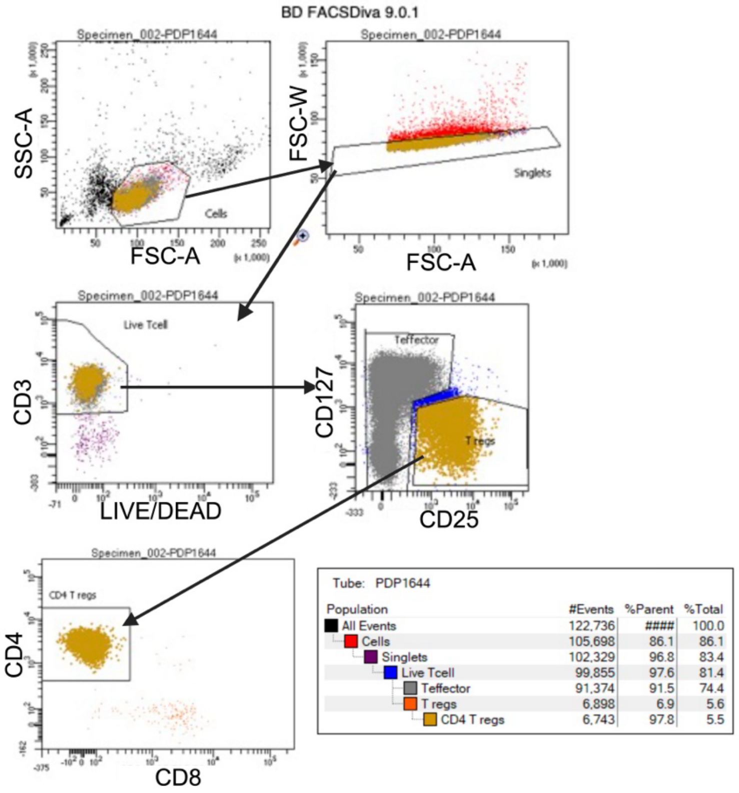


**S2.** ***FACS sort gating for suppression assays.*** *Gating strategy: lymphocytes were gated on SSC-A against FSC-A. Within this population, singlets were gated using FSC-W against FSC-A. Live T cells were identified as singlets that were CD3+LIVE/DEAD-. Within this live T cell population, T effectors were gated as CD25low and Tregs as CD127lowCD25high. The Treg population was further subdivided into CD4 Tregs and only CD4 Tregs were sorted.*

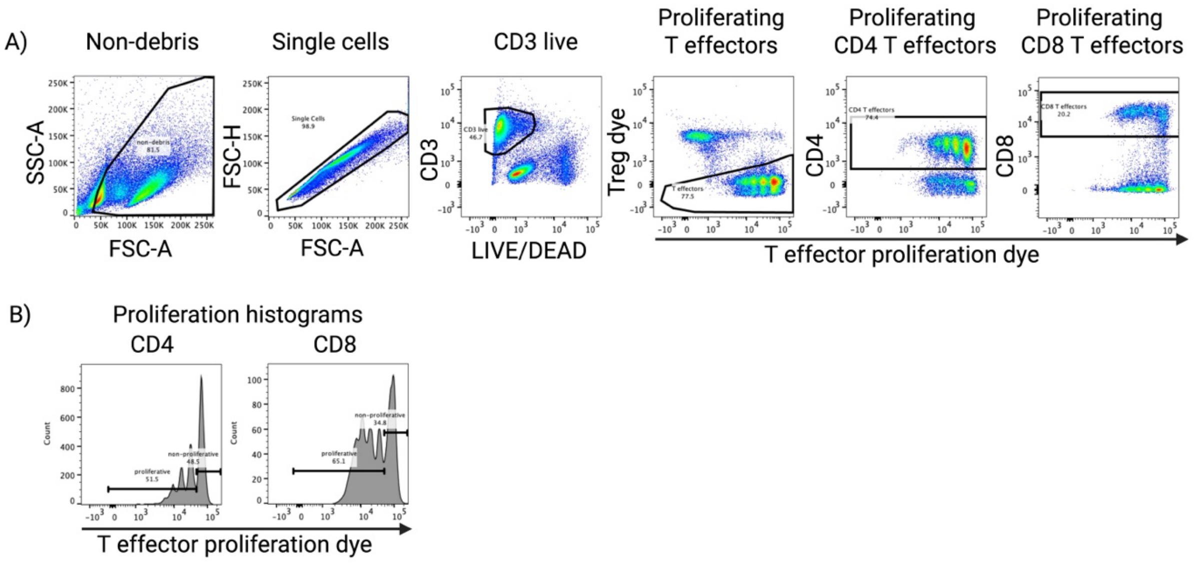


**S3.** ***Gating strategy used for analysis of suppression assays. A)*** *Dot plots showing gating of proliferating T effectors. Non-debris was gated on SSC-A vs FSC-A. Single cells were gated*

*on FSC-H vs FSC-A. Live T cells were gated as CD3+LIVE/DEAD-. Proliferating T effectors were identified as Treg dye (670)- and varying intensities of T effector proliferation dye (450). CD4 T effectors were CD4+ vs T effector proliferation dye and CD8 T effectors were CD8+ vs T effector proliferation dye.* ***B)*** *Histograms showing how percentage proliferation was calculated using proliferation dye (x-axis). A non-proliferative gate was drawn using the unstimulated T effector well and this gate was then applied to all wells. Proliferative effectors were defined as any cells to the left of this region.*

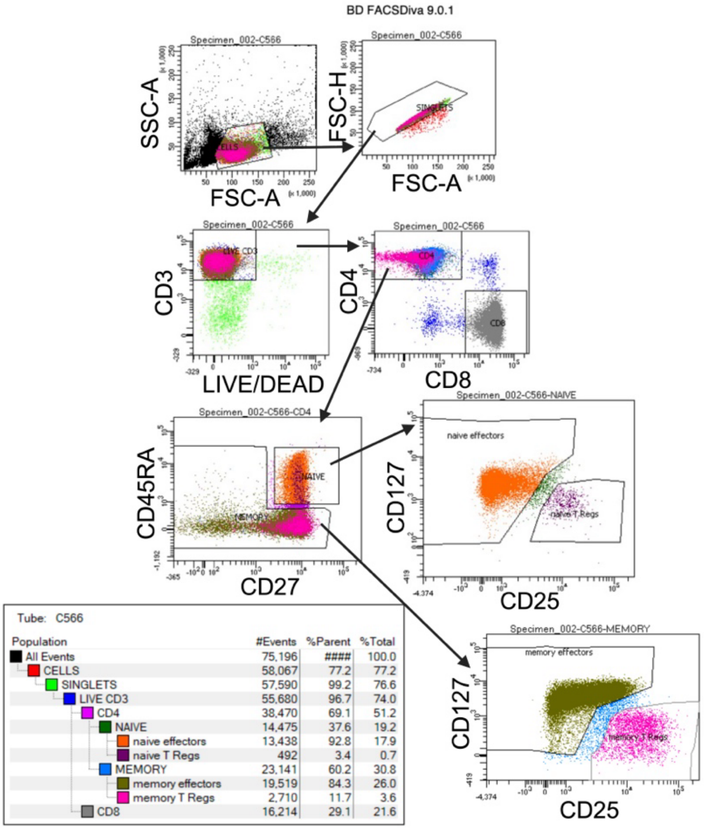


| Population | #Events | %Parent | %Total |
|---|---|---|---|
| All Events | 75,196 | #### | 100.0 |
| CELLS | 58,067 | 77.2 | 77.2 |
| SINGLETS | 57,590 | 99.2 | 76.6 |
| LIVE CD3 | 55,680 | 96.7 | 74.0 |
| CD4 | 38,470 | 69.1 | 51.2 |
| NAIVE | 14,475 | 37.6 | 19.2 |
| naive effectors | 13,438 | 92.8 | 17.9 |
| naive T Regs | 492 | 3.4 | 0.7 |
| MEMORY | 23,141 | 60.2 | 30.8 |
| memory effectors | 19,519 | 84.3 | 26.0 |
| memory T Regs | 2,710 | 11.7 | 3.6 |
| CD8 | 16,214 | 29.1 | 21.6 |

**S4.** ***Gating strategy for the FACS sort to isolate naive and memory Tregs for TSDR demethylation analysis.***

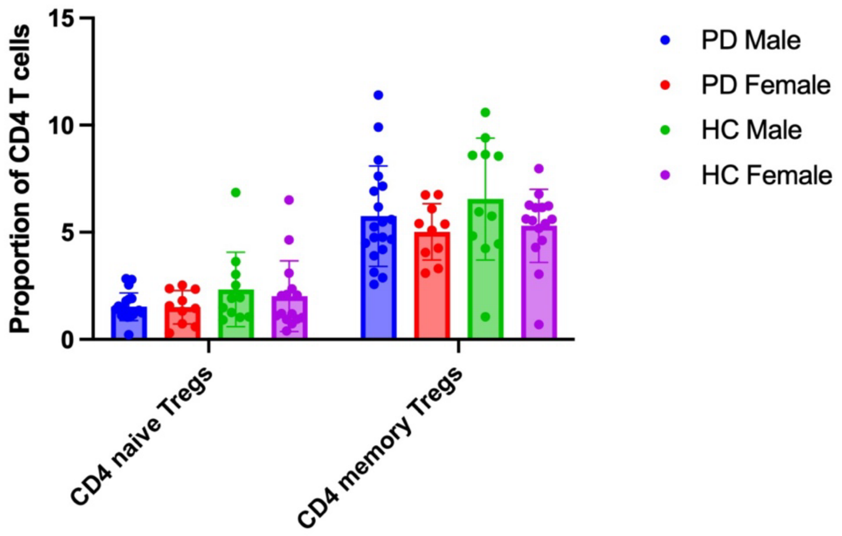


**S5. *Stratifying naive and memory Tregs as a proportion of CD4 T cells by sex.*** *Graph shows mean ±SD. PD, n = 19M, 10F; HC, n = 11M, 15F.*

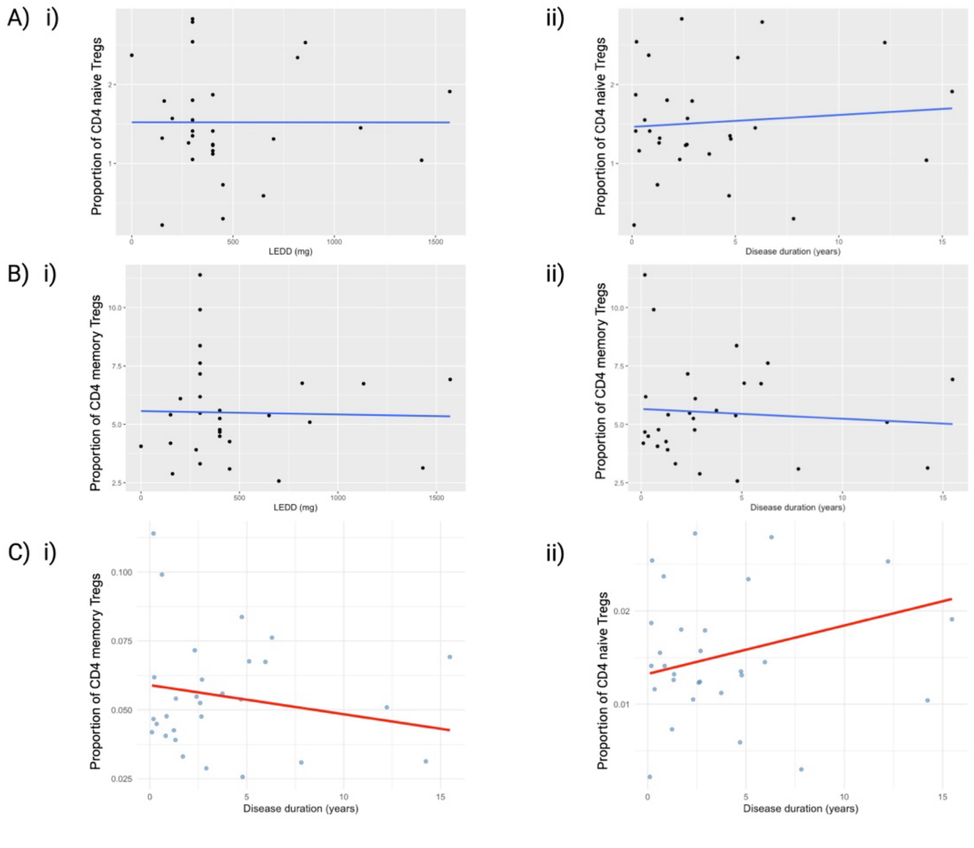


**S6. *Linear models for correlations between disease duration, LEDD and Treg proportions.*** ***A)*** *Proportion of CD4 naive Tregs against* ***i)*** *LEDD* ***ii)*** *disease duration.* ***B)*** *Proportion of CD4 memory Tregs against* ***i)*** *LEDD* ***ii)*** *disease duration.* ***C)*** *Mixed effects model for correlation between proportion of* ***i)*** *CD4 memory Tregs* ***ii)*** *CD4 naive Tregs against disease duration, correcting for LEDD. PD, n = 29.*

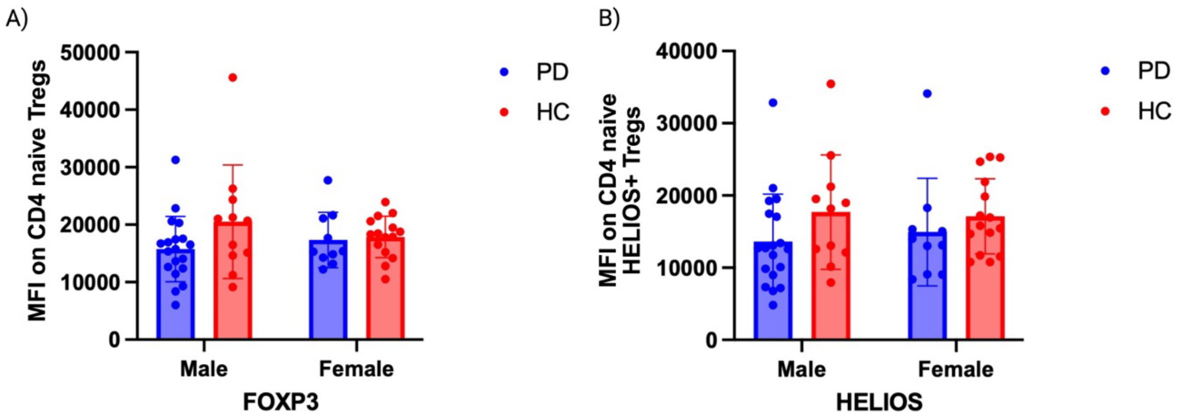


**S7.** ***Impact of sex on FOXP3 and HELIOS MFI. A)*** *CD4 naive Treg FOXP3 MFI of PD and HC males and females.* ***B)*** *CD4 naive HELIOS+ Treg MFI of PD and HC males and females. Graphs show mean ±SD. PD, n = 19M, 10F; HC, n = 11M, 15F.*
